# Gpnmb inhibits oligodendrocyte differentiation of adult neural stem cells by amplifying TGFβ1 signaling

**DOI:** 10.1101/2021.08.13.456269

**Authors:** Daniel Z. Radecki, Albert R. Wang, Abigail S. Johnson, Christian A. Overman, Madison M. Thatcher, Gopal Iyer, Jayshree Samanta

## Abstract

Gli1 expressing neural stem cells, in the subventricular zone of the adult mammalian brain, respond to demyelination injury by differentiating into oligodendrocytes. We have identified Gpnmb as a novel regulator of oligodendrogenesis in Gli1 neural stem cells, whose expression is induced by TGFβ1 signaling via Gli1, in response to a demyelinating injury. Upregulation of Gpnmb further activates the TGFβ1 pathway by increasing the expression of the TGFβ1 binding receptor subunit, TGFβR2. Thus the TGFβ1→Gli1→Gpnmb→TGFβR2 signaling pathway forms a feed forward loop for sustained activation of TGFβ1 signaling in Gli1 neural stem cells, resulting in inhibition of their differentiation into mature oligodendrocytes following demyelination.

## INTRODUCTION

The adult subventricular zone (SVZ) consists of quiescent neural stem cells that are multipotent, divide slowly and self-renew (Doetsch et al., 1999). There is considerable heterogeneity amongst these cells, indicated by their expression of specific transcription factors and generation of distinct cells in homeostasis and injury (Chaker et al., 2016). One of the subsets residing in the ventral SVZ accounts for 25% of the quiescent neural stem cells, expresses the transcription factor Gli1, and generates interneurons in the olfactory bulb and astrocytes in the healthy mouse forebrain (Ahn and Joyner, 2005). These ventral neural stem cells (vNSCs) are also present in the human SVZ (Samanta et al., 2015). Remarkably, the Gli1 vNSCs can be recruited to the white matter where they differentiate into myelinating oligodendrocytes in response to demyelination but not in the healthy adult brain (Samanta et al., 2015; Sanchez and Armstrong, 2018; Sanchez et al., 2018). Inhibition of Gli1 in these cells further enhances their remyelination potential and is neuroprotective in mouse models of multiple sclerosis (MS) (Samanta et al., 2015). In this study we analyzed the transcriptome of vNSCs to discover that repression of TGFβ1 signaling pathway along with downregulation of glycoprotein non-metastatic melanoma b (Gpnmb) are accompanied by loss of Gli1 in these cells, following demyelination.

The TGFβ family influences proliferation, migration and differentiation of NSCs, along with modulating the immune system (David and Massague, 2018). Indeed, TGFβ ligands including TGFβ1 as well as the receptor subunits TGFβR1 and TGFβR2 are highly expressed by reactive astrocytes and microglia in chronic MS lesions (De Groot et al., 1999). TGFβ1 is also upregulated in the aging brain where remyelination is limited (Nicaise et al., 2019; Pasinetti et al., 1999). In rodents, demyelinating lesions show higher TGFβ1 expression (Hinks and Franklin, 2000) and transgenic overexpression of TGFβ1 in the brain results in earlier onset and more severe disease in the EAE model suggesting a negative role in remyelination (Wyss-Coray et al., 1997). On the contrary, genetic knockout of TGFβ1 in the developing embryo results in focal demyelination in the brain suggesting a positive effect on developmental myelination (Makwana et al., 2007). Thus, the actions of TGFβ1 signaling are dependent on the timing, concentration and cell specific modulators of the pathway. TGFβ1 signals by binding to the receptor TGFβR2 leading to recruitment and phosphorylation of TGFβR1 receptor thus activating its kinase. Activated TGFβR1 phosphorylates Smad2/3, which form a complex with Smad4 and translocate to the nucleus for regulation of transcription (Massague, 1987; Wrana et al., 1994).

In this study, we identified a novel regulator of remyelination - Gpnmb, via its role in activation of the TGFβ1 pathway by upregulating TGFβR2 expression in vNSCs. Gpnmb is a single-pass transmembrane glycoprotein with a C-terminus cytoplasmic domain and an N-terminus extracellular domain. The human and mouse genes are very similar with a high degree of conservation in the chromosomal region around Gpnmb in both genomes (Owen et al., 2003). Gpnmb is upregulated in many neurodegenerative diseases featuring demyelination such as in amyotrophic lateral sclerosis, Alzheimer’s disease, Parkinson’s disease and MS (Hendrickx et al., 2017; Hüttenrauch et al., 2018; Moloney et al., 2018; Satoh et al., 2019; Tanaka et al., 2012). In MS brains, Gpnmb is highly expressed in chronic active lesions, which are associated with more aggressive disease and declining remyelination (Hendrickx et al., 2017).

In this study, we have used a Gpnmb-LacZ knock-in mouse to not only characterize the previously unknown expression of Gpnmb in the adult forebrain but also to elucidate its role in regulation of remyelination by vNSCs. Our results indicate that demyelination induces TGFβ1, which acts via Gli1 to upregulate Gpnmb in vNSCs; overexpression of Gpnmb further induces expression of the TGFβR2 receptor thus enhancing the activation of the TGFβ1 pathway, leading to the reduced generation of mature OLs from vNSCs.

## RESULTS

### Identification of Gpnmb as a novel inhibitor of oligodendrogenesis in ventral neural stem cells

Our previous studies have shown that loss of Gli1 in vNSCs of the adult SVZ, results in enhanced recruitment and regeneration of myelinating oligodendrocytes in response to demyelination of the white matter corpus callosum (CC) (Radecki et al., 2020; Samanta et al., 2015). To determine the molecular mechanisms involved, we compared the transcriptomes of vNSCs isolated from the healthy and demyelinated brains of the knock-in Gli1^CreER/Wt^ (*Gli1^HET^*) mice with one copy of Gli1, and Gli1^CreER/CreER^ (*Gli1^NULL^*) mice with loss of Gli1 expression (Samanta, 2021). We probed the top 100 differentially expressed genes in the *Gli1^HET^* vs. *Gli1^NULL^* vNSCs following demyelination, for genes shared amongst three or more GO terms related to cell survival, migration and differentiation. Through this analysis, we identified *Gpnmb*, which was also the most significantly downregulated gene, with 7.9 fold lower expression, in *Gli1^NULL^* vNSCs, suggestive of its role as a negative regulator of remyelination (Fig.1A). Notably, *Gpnmb* was upregulated following demyelination in *Gli1^HET^* vNSCs but not in the *Gli1^NULL^* vNSCs (Fig.1A), further suggesting that lack of *Gpnmb* induction in *Gli1^NULL^* vNSCs may be beneficial for remyelination.

**Figure 1.**
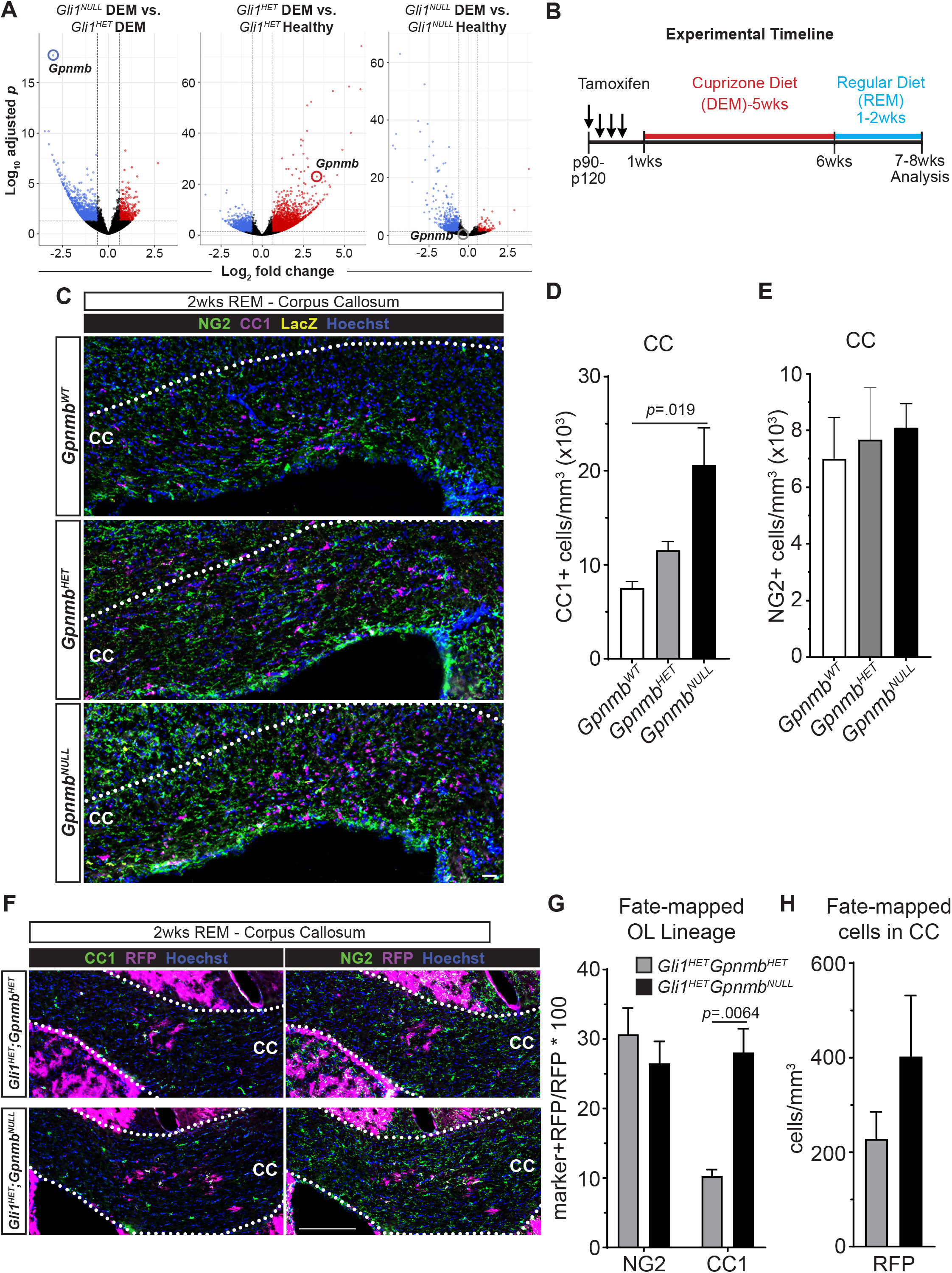
Loss of *Gpnmb* increases differentiation of Gli1 vNSCs into oligodendrocytes following demyelination. (A) RNAseq comparing *Gli1^NULL^* to *Gli1^HET^* vNSCs, identified *Gpnmb* as the most significantly downregulated gene in *Gli1^NULL^* vNSCs following demyelination (Left). *Gpnmb* is significantly upregulated in *Gli1^HET^* vNSCs on demyelination as compared to vNSCs from healthy mice (middle). *Gpnmb* is not induced upon demyelination in *Gli1^NULL^* vNSCs (right). (B) Schematic of the experimental timeline for Cuprizone induced demyelination. (C) Immunofluorescent images of the CC at 2 weeks of recovery from cuprizone diet showing NG2+ OPCs (green), CC1+ OLs (magenta) and LacZ+ Gpnmb expressing cells (yellow) in the CC. Scale = 30µm (D) Quantification shows a significant increase in CC1+ OLs in the *Gpnmb^NULL^* CC compared to *Gpnmb^WT^* CC at 2 weeks of remyelination. n=3, data=mean±SEM, one-way ANOVA with post-hoc t-tests. (E) Quantification for NG2+ OPCs in the CC showed no change at 2 wks of remyelination. n=4, 1-way ANOVA with post-hoc t-test. (F) Immunofluorescent images of the *Gli1^HET^;Gpnmb^HET^* and *Gli1^HET^;Gpnmb^NULL^* CC at 2 weeks of remyelination showing RFP+ fate-mapped Gli1 vNSCs (magenta) co-expressing CC1 (green) and NG2 (green). Nuclei are counterstained with Hoechst. Scale = 25µm. (G) Quantification of percentage of RFP+ Gli1 fate-mapped vNSCs co-expressing NG2 and CC1, shows a significant increase in CC1+ OLs in *Gli1^HET^;Gpnmb^NULL^* CC compared to *Gli1^HET^;Gpnmb^HET^* CC at 2 weeks of remyelination. n=3, unpaired t-tests within groups. (H) Quantification of total number of RFP+ Gli1 fate-mapped cells in the *Gli1^HET^;Gpnmb^HET^* and *Gli1^HET^;Gpnmb^NULL^* CC at 2 weeks of remyelination. DEM-demyelination, REM-remyelination, OPC-oligodendrocyte progenitor cells, OL-oligodendrocytes, CC-corpus callosum

To determine if *Gpnmb* plays a role in remyelination, we examined the consequences of genetic loss of *Gpnmb* using a β-Galactosidase knock-in Gpnmb^LacZ^ mouse (Table S1), thus allowing us to identify the cells with an active *Gpnmb* promoter even after complete loss of *Gpnmb* expression in *Gpnmb^NULL^* (Gpnmb^LacZ/LacZ^) mice. Global loss of *Gpnmb* did not result in any overt phenotype and the mice were healthy and fertile with a normal lifespan. We induced demyelination in the *Gpnmb^NULL^* (Gpnmb^LacZ/LacZ^) and *Gpnmb^HET^* (Gpnmb^LacZ/Wt^) mice by feeding cuprizone diet for 5 weeks and analyzed the brains at 2 weeks of recovery from the diet (Fig.1B). In this model, cuprizone diet results in peak demyelination at 5-6 weeks and returning the mice to regular chow allows remyelination to occur spontaneously (Matsushima and Morell, 2001). While the extent of demyelination was similar in *Gpnmb^NULL^* and *Gpnmb^WT^* mice (Fig. S1A), the number of CC1-expressing mature oligodendrocytes in the CC of *Gpnmb^NULL^* mice was 2.7±0.1 fold higher than in the *Gpnmb^WT^* and 1.8±0.4 fold higher than *Gpnmb^HET^* mice (Fig.1 C,D). However, the number of NG2+ oligodendrocyte progenitor cells (OPCs) did not change upon loss of Gpnmb (Fig.1C,E). Taken together these results indicate that *Gpnmb* inhibits oligodendroglial differentiation in the white matter following demyelination.

To determine if the increase in OLs in the CC following demyelination was due to an increase in recruitment and/or differentiation of vNSCs, we traced the lineage of Gli1 vNSCs by genetically fate-mapping the cells in the *Gpnmb^NULL^* mice at 2 weeks of recovery from cuprizone diet, using G*li1^HET^;Gpnmb^NULL^* mice (Table S1). Consistently, we found a significant 2.7±0.1 fold increase in the proportion of CC1+RFP double positive cells derived from fate-mapped vNSCs in the *Gli1^HET^;Gpnmb^NULL^* CC compared to the *Gli1^HET^;Gpnmb^HET^* CC, indicating an increase in generation of mature OLs from the Gli1 vNSCs (Fig.1F,G). However, there was no change in the number of NG2+RFP double positive OPCs or GFAP+RFP double positive astrocytes derived from fate-mapped vNSCs between the *Gli1^HET^;Gpnmb^HET^* and *Gli1^HET^;Gpnmb^NULL^* mice (Fig.1F,G; Fig.S1B). In addition, the progeny of Gli1 vNSC cells did not differentiate into NeuN+ neurons or Iba1+ microglia (Fig.S1B). Thus, the Gli1 vNSCs generate more OLs without affecting their differentiation into OPCs, astrocytes, neurons or microglia. To examine if the increase in OLs was in part due to an increase in the number of Gli1 vNSCs recruited to the CC, we quantified the total number of RFP+ fate-mapped Gli1 vNSCs in the CC, but did not observe a significant change in the *Gli1^HET^;Gpnmb^NULL^* mice compared to the *Gli1^HET^;Gpnmb^HET^* mice (Fig.1F,H). In addition, there was no change in the number of fate-mapped RFP+ vNSCs with an active Gpnmb promoter (LacZ+), between the *Gli1^HET^;Gpnmb^HET^* and *Gli1^HET^;Gpnmb^NULL^* mice (Fig.S1C-D). These data indicate that loss of Gpnmb increases the differentiation of cells derived from Gli1 vNSC into OLs, in the white matter following demyelination.

### Expression of Gpnmb is limited to NSCs and glial cells in the forebrain

To characterize the expression of Gpnmb in the quiescent NSCs, we first quantified the number of LacZ+ cells co-expressing GFAP in the entire SVZ at 1 week of recovery from cuprizone diet as compared to the healthy SVZ in *Gpnmb^HET^* mice, but did not observe any difference between the healthy and remyelinating SVZ (Fig.2A-B). In addition there were no differences in the proportion of LacZ+ cells co-expressing markers of OLs (CC1), OPCs (PGFRα), neurons (NeuN) and microglia (Iba1) in the healthy vs. remyelinating SVZ (Fig.2B). To examine the expression of Gpnmb specifically in Gli1 vNSCs, we quantified the RFP+ fate-mapped Gli1 cells co-expressing LacZ in the *Gli1^HET^;Gpnmb^HET^* mice. There was a significant 1.5±0.1 fold increase in the proportion of RFP+LacZ double positive cells in the SVZ, at 1 week of recovery from cuprizone diet (45.9±2.6%) compared to the healthy SVZ (31.3±3.9%) (Fig.2C-D), consistent with the RNAseq data showing increase in Gpnmb expression upon demyelination in the *Gli1^HET^* NSCs.

**Figure 2.**
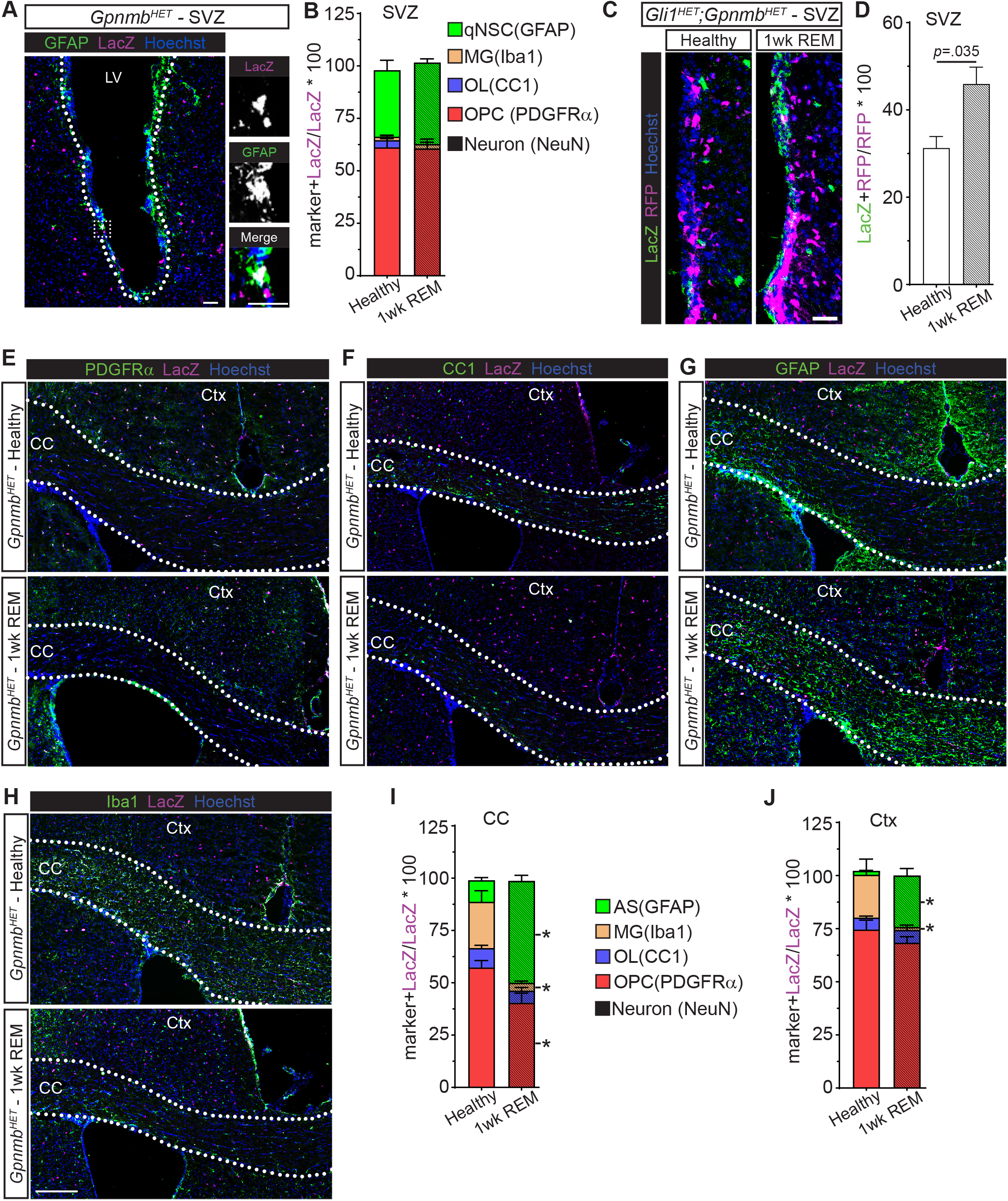
Expression of Gpnmb is limited to NSCs and glial cells in the forebrain. (A) Immunofluorescent image showing co-localization of LacZ (magenta) and GFAP (green) in qNSCs (inset) in the SVZ (large dotted outline) of healthy *Gpnmb^HET^* mice. Scale=25□m. (B) Quantification of the percentage of LacZ+ cells co-expressing markers of qNSCs (GFAP), microglia (Iba1), oligodendrocytes (CC1), OPCs (PDGFRα) and neurons (NeuN), in the SVZ of healthy *Gpnmb^HET^* mice or at 1 week of recovery from cuprizone diet (REM). No co-localization with NeuN+ neurons was detected. n=3, 2-way ANOVA with post-hoc t-tests within groups. (C) Immunofluorescent images showing co-localization of LacZ (green) and RFP (magenta) in the healthy CC and after 1 week of remyelination in *Gli1^HET^;Gpnmb^HET^* mice. Scale=25µm (D) Quantification of (C) showing an increase in the proportion of RFP+ Gli1 vNSCs expressing LacZ at 1 week of recovery from cuprizone diet (REM) compared to healthy SVZ. n=3, unpaired t-test. (E) Immunofluorescent image from *Gpnmb^HET^* mice showing PDGFRα+ OPCs and LacZ+ cells in the healthy forebrain and after 1 week of remyelination. Scale=100µm (F) Immunofluorescent image from *Gpnmb^HET^* mice showing CC1+ OLs and LacZ+ cells in the healthy forebrain and after 1 week of remyelination. Scale=100µm (G) Immunofluorescent images from *Gpnmb^HET^* mice showing GFAP+ astrocytes and LacZ+ cells in the healthy forebrain and after 1 week of remyelination. Scale=100µm (H) Immunofluorescent images from *Gpnmb^HET^* mice showing Iba1+ microglia and LacZ+ cells in the healthy forebrain and after 1 week of remyelination. Scale=100µm (I) Quantification of proportion of LacZ+ cells co-expressing cell-type specific markers in the healthy CC (solid bars) and at 1wk REM (thatched bars). There is a significant increase in the proportion of LacZ+ astrocytes and a decrease in microglia and OPCs. No co-localization with NeuN+ neurons was detected. n=3, 2-way ANOVA with post-hoc t-tests within groups. (J) Quantification of proportion of LacZ+ cells co-expressing cell-type specific markers in the healthy cortex (solid bars) and at 1wk REM (thatched bars).There is a significant increase in the proportion of LacZ+ astrocytes and a decrease in microglia. No co-localization with NeuN+ neurons was detected. n=3, 2-way ANOVA with post-hoc t-tests within groups. \**p*<.0001. Quiescent neural stem cell (qNSC), ventral neural stem cell (vNSC), Lateral ventricle (LV), corpus callosum (CC), cortex (Ctx), subventricular zone (SVZ), oligodendrocyte precursor cell (OPC), oligodendrocyte (OL), astrocyte (AS), microglia (MG)

Loss of Gpnmb can potentially enhance remyelination due to its effects on OPCs and OLs. In addition, several RNAseq studies have indicated that *Gpnmb* is expressed in oligodendroglial and microglial cells in the mouse and human brain (Srinivasan et al., 2020; Zhang et al., 2014; Zhang et al., 2016). Therefore, we characterized the expression of Gpnmb in the CC and cortex of the healthy adult *Gpnmb^HET^* forebrain and following 1 week of recovery from cuprizone induced demyelination. In the healthy CC, the Gpnmb expressing cells consisted of GFAP+ astrocytes (10.3±1.3%), Iba1+ microglia (22.1±2.6%), PDGFRα+ OPCs (57.3±3.3%) and CC1+ OLs (9.3±1.3%) (Fig.2 E-I). However, at 1wk of recovery from cuprizone diet, Gpnmb expressing GFAP+ astrocytes increased (4.7±0.3 fold) with a concomitant significant reduction in Iba1+ microglia (0.17±0.02 fold) and PDGFRα+ OPCs (0.7±0.1 fold) (Fig.2 E-I).

In the healthy cortex, the Gpnmb expressing cells were comprised of 1.8±0.2% GFAP+ astrocytes, 20.2±3.7% Iba1+ microglia, 74.6±4.4% PDGFRα+ OPCs and 5.6±0.6% CC1+ OLs (Fig.2 E-J). Compared to the healthy CC, the cortex had 1.3±0.1 fold higher number of Gpnmb+ OPCs and a 0.18±0.02 fold lower number of Gpnmb+ astrocytes. Similar to the CC, Gpnmb expression significantly increased by 13.2±1.8 fold in astrocytes and decreased by 0.07±0.02 fold in microglia, in the cortex at 1 week of recovery from cuprizone diet (Fig.2 E-J). Notably, Gpnmb was not expressed in neurons in the healthy or remyelinating SVZ, CC, and cortex.

Thus, Gpnmb is expressed in astroglial, oligodendroglial and microglial cells along with NSCs in the healthy forebrain and its expression is dynamically regulated by demyelinating injury.

### Gpnmb is necessary for inhibition of oligodendrogenesis by TGFβ1

By performing an Ingenuity and Kegg pathway analyses of the RNAseq data (Samanta, 2021), we identified the TGFβ1 pathway as significantly activated in *Gli1^HET^* vNSCs as compared to *Gli1^NULL^* vNSCs following demyelination (Fig.3A). To confirm the source of TGFβ1 in the adult brain, we compared its mRNA expression in the CC, cortex and SVZ of healthy vs demyelinated brain. After 3 weeks of cuprizone diet, the timepoint which coincides with the initiation of demyelination, *Tgfb1* expression showed a significant 2.2±0.4 fold increase in the CC, but remained unchanged in the cortex and SVZ (Fig.3B). This suggests that TGFβ1 is secreted from the lesion in response to a demyelinating injury. To examine if TGFβ1 can regulate Gpnmb expression, we harvested vNSCs from the SVZ of adult *Gli1^HET^* and *Gli1^NULL^* mice and grew them as neurospheres *in vitro*, followed by differentiation of the dissociated neurospheres in adherent cultures for 14 days (Fig.3C). Treatment of vNSCs with TGFβ1 (10ng/mL), significantly increased *Gpnmb* mRNA levels by 4.3±1.2 fold in *Gli1^HET^* vNSCs, but not in *Gli1^NULL^* vNSCs (Fig. 3D). Moreover this effect on Gpnmb upregulation was very specific to TGFβ1, since treatment with other growth factors like Shh (100ng/mL) (Galvin et al., 2008) and BMP-4 (20ng/mL) (Bond et al., 2014; Samanta et al., 2007) failed to change Gpnmb expression in both *Gli1^HET^* and *Gli1^NULL^* vNSCs (Fig.3D). These results suggest that TGFβ1 acts via Gli1 to upregulate Gpnmb expression in *Gli1^HET^* vNSCs, and consequently fails to induce Gpnmb in *Gli1^NULL^* vNSCs.

**Figure 3.**
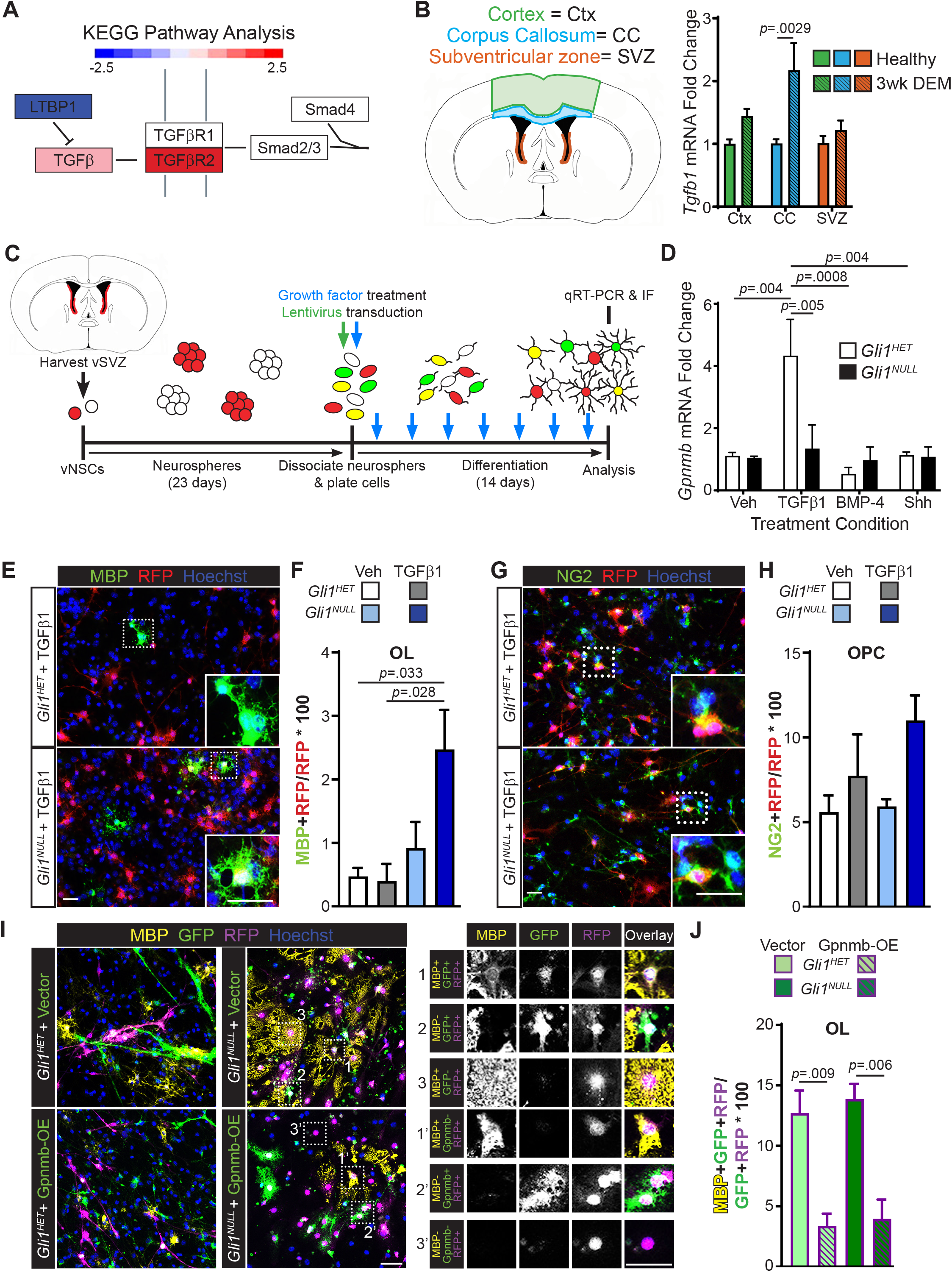
TGFβ1 inhibits oligodendrogenesis by induction of *Gpnmb* expression. (A) KEGG pathway analysis of the RNAseq data showing activation of the TGFβ pathway in *Gli1^HET^* vNSCs compared to *Gli1^NULL^* vNSCs upon demyelination. (B) Schematic showing the CNS regions dissected for mRNA extraction (left). QRT-PCR for TGFβ1 expression in the CC, Ctx and SVZ tissue dissected from adult wildtype C57bl/6 mice fed with regular diet (Healthy) or cuprizone diet (DEM) for 3 weeks shows significant upregulation in the CC at 3wks demyelination (right). n=3, unpaired t-test within dissection regions. (C) Experimental timeline for establishing *in vitro* culture of vNSCs. (D) QRT-PCR for *Gpnmb* expression in vNSCs harvested from *Gli1^HET^* and *Gli1^NULL^* mice and treated with TGFβ1, BMP-4 and Shh *in vitro*, shows a significant increase in Gpnmb expression only in *Gli1^HET^ v*NSCs following TGFβ1 treatment. n=3, 2-way ANOVA with post-hoc t-tests between all conditions. (E) Immunofluorescent images from *Gli1^HET^* and *Gli1^NULL^* vNSCs following treatment with TGFβ1 showing RFP+ cells derived from Gli1 vNSC and MBP+ OLs (insets). Scale = 25µm (F) Quantification of (E) showing an increase in MBP+ OLs from *Gli1^NULL^* vNSCs upon TGFβ1 treatment, compared to *Gli1^HET^* vNSCs. n=3, 1-way ANOVA with post-hoc t-tests between all groups. (G) Immunofluorescent images from *Gli1^HET^* and *Gli1^NULL^* vNSCs following treatment with TGFβ1 showing NG2+ OPCs co-expressing RFP (insets). Scale = 25µm (H) Quantification of (G) shows no difference in RFP+ OPCs generated from *Gli1^HET^* and *Gli1^NULL^* vNSCs with TGFβ1 treatment. n=3, 1-way ANOVA with post-hoc t-tests between all groups. (I) Immunofluorescent images from *Gli1^HET^* and *Gli1^NULL^* vNSCs following overexpression of Gpnmb using lentiviruses, showing GFP+ cells infected with the lentivirus (green), RFP+ cells derived from Gli1 vNSCs (magenta), and MBP+ OLs (yellow). Insets highlight cells showing co-localization of the markers- (1) GFP+RFP+MBP+ cell, (2) GFP+RFP+MBP-cell, (3) GFP-RFP+MBP+ cell, (1’) GFP-RFP-MBP+ cell, (2’) GFP+RFP+MBP-cell, (3’) GFP-RFP+MBP+ cell. Scale = 25µm (J) Quantification of the percentage of GFP+RFP+ OLs after 14 days of differentiation of vNSCs *in vitro,* showing significantly fewer OLs generated from both *Gli1^HET^* and *Gli1^NULL^* vNSCs upon *Gpnmb* overexpression (Gpnmb-OE) n=3, 1-way ANOVA with post-hoc t-tests between all groups.

To examine the effect of TGFβ1 on differentiation of vNSCs, we quantified the proportion of RFP+ cells co-expressing markers of OLs, OPCs, or astrocytes following TGFβ1 treatment. Although TGFβ1 did not alter the generation of OLs from *Gli1^HET^* vNSCs, it significantly increased the differentiation of *Gli1^NULL^* vNSCs into OLs by 5.2±1.3 fold (Fig.3 E-F) and decreased their differentiation into astrocytes by 0.5±0.1 fold (Fig.S2 A-B). Consistent with previous results there was no change in the differentiation of *Gli1^HET^* and *Gli1^NULL^* vNSCs into OPCs (Fig.3G,H). Thus, TGFβ1 treatment upregulates Gpnmb but does not affect the generation of OLs from *Gli1^HET^* vNSCs. In contrast, it increases the differentiation of *Gli1^NULL^* vNSCs into OLs, without affecting Gpnmb expression. This suggests that upregulation of Gpnmb by TGFβ1, inhibits the differentiation of vNSCs into OLs.

To directly examine the inhibitory role of Gpnmb in oligodendrogenesis, we overexpressed *Gpnmb* with lentiviral transduction of the Gpnmb cDNA fused with GFP (Gpnmb-OE), which significantly increased Gpnmb mRNA levels by 402±141 fold in vNSCs (Fig.S3A). The efficiency of lentiviral infection was similar in both *Gli1^HET^* and *Gli1^NULL^* vNSCs (Fig.S3B). We then quantified the number of OLs, OPCs and astrocytes generated from *Gli1^HET^* and *Gli1^NULL^* vNSCs overexpressing Gpnmb i.e. proportion of RFP+GFP double positive cells co-expressing the cell type specific markers. Gpnmb overexpression significantly decreased the generation of OLs by 3.8±0.6 fold in *Gli1^HET^* and 3.5±0.3 fold in *Gli1^NULL^* vNSCs (Fig.3 I,J). However it did not alter the differentiation of vNSCs into OPCs (Fig.S3 C,D) and astrocytes (Fig.S3 E,F).

Together, these results indicate that demyelination increases TGFβ1 expression in the lesion, which induces *Gpnmb* expression via Gli1, leading to reduction in generation of mature OLs from vNSCs.

### Gpnmb inhibits oligodendrogenesis by inducing TGFβR2 expression

Since the RNAseq data (Samanta, 2021) showed significantly higher expression of the ligand binding subunit of TGFβ receptor, TGFβR2, in the *Gli1^HET^* vNSCs compared to the *Gli1^NULL^* vNSCs upon demyelination (Fig.3 A), it suggested that Gpnmb can regulate the TGFβ1 pathway. Consistently, overexpression of *Gpnmb* in wildtype vNSCs *in vitro,* resulted in a significant upregulation of *Tgfbr2* by 2.2±0.5 fold but did not change *Tgfbr1* subunit mRNA expression (Fig.4A), indicating a potential feed forward loop consisting of TGFβ1→Gli1→Gpnmb→TGFβR2. To confirm this, we examined *Tgfbr2* expression in the SVZ of *Gpnmb^HET^* and *Gpnmb^NULL^* mice. Consistent with the data from vNSCs, *in vivo* loss of *Gpnmb* also resulted in a significant reduction in *Tgfbr2* expression by 0.54±0.05 fold without altering *Tgfbr1* expression and this effect was not due to loss of *Tgfb1* expression in *Gpnmb^NULL^* mice (Fig.4B). Thus TGFβ1 induces Gpnmb via Gli1, which in turn induces TGFβR2.

**Figure 4.**
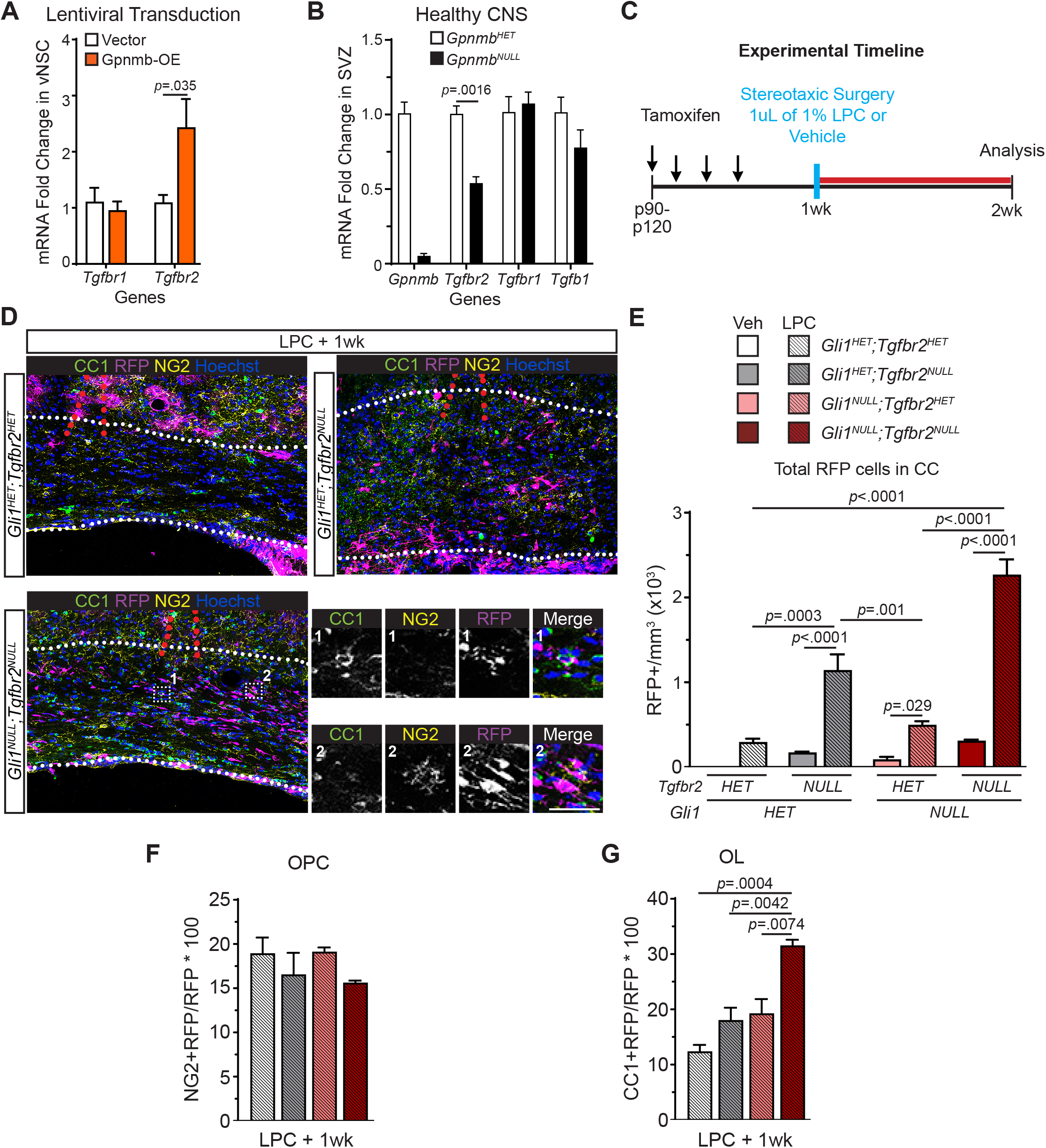
Conditional ablation of TGFβR2 in vNSCs increases their recruitment and oligodendrogenesis in Gli1^NULL^ mice following demyelination. (A) QRT-PCR for the TGFβ1 receptor subunits, Tgfbr1 and Tgfbr2, with overexpression of Gpnmb (orange) or empty vector (white) in vNSCs harvested from adult wildtype C57bl/6 mice, shows significant increase in the ligand binding *Tgfbr2* subunit. n=3, one-way ANOVA with post-hoc t-tests within groups. (B) QRT-PCR for expression of Gpnmb, TGFβ1 and TGFβ1 receptor subunits in the *Gpnmb^HET^* and *Gpnmb^NULL^* SVZ, shows a significant decrease in *Tgfbr2* expression. n=4, data=mean±SEM, unpaired t-test within gene groups. (C) Schematic of the experimental timeline for lysophosphatidyl choline (LPC) induced demyelination. (D) Immunofluorescent images showing co-localization of RFP (magenta) with CC1 (green) or NG2 (yellow) in the CC of *Gli1^HET^Tgfbr2^HET^*, *Gli1^HET^Tgfbr2^NULL^,* and G*li1^NULL^Tgfbr2^NULL^* mice. Scale = 30µm (E) Quantification of (D) shows increase in number of RFP+ cells derived from vNSCs in the corpus callosum (CC) of both *Gli1^HET^* and *Gli1^NULL^* mice upon loss of *Tgfbr2*. n=3, 2-way ANOVA with post-hoc t-tests between all groups. (F) Quantification of percentage of RFP+ OPCs in the corpus callosum (CC) shows no change with loss of *Tgfbr2* in *Gli1^HET^* and *Gli1^NULL^* mice following demyelination n=3, 1-way ANOVA with post-hoc t-tests. (G) Quantification of percentage of RFP+ OLs in the corpus callosum (CC) shows an increase with loss of *Tgfbr2* in *Gli1^NULL^* mice following demyelination n=3, 1-way ANOVA with post-hoc t-tests.

Next, we wanted to confirm whether loss of TGFβR2 was sufficient to alter vNSC recruitment or differentiation *in vivo.* We conditionally knocked out *Tgfbr2* from *Gli1^HET^* and *Gli1^NULL^* vNSCs and examined the fate of their progeny after inducing demyelination by stereotactic injection of Lysophosphatidyl choline (LPC) (Plemel et al., 2018) (Fig.4 C). We confirmed the ablation of TGFβ1 signaling in Gli1 vNSCs, by the absence of pSmad3, a downstream effector of TGFβ1 signaling pathway (Massague, 1987; Wrana et al., 1994) (Fig.S4A). In addition, we confirmed that the extent of demyelination induced by LPC injection was similar in both *Tgfbr2^HET^* and *Tgfbr2^NULL^* mice (Fig.S4B).

We then analyzed the recruitment of the *Gli1^HET^* and *Gli1^NULL^* vNSCs to the CC, 1 week after LPC or vehicle injection. The total number of RFP+ cells in the lesion, was significantly higher by 6.6±1.1 fold in *Gli1^HET^; Tgfbr2^NULL^*, 5.4±0.4 fold in *Gli1^NULL^;Tgfbr2^HET^* and 7.3±0.6 fold in *Gli1^NULL^;Tgfbr2^NULL^* mice as compared to the respective vehicle injected controls (Fig.4D,E). Thus, while complete loss of *Tgfbr2* increased the recruitment of both *Gli1^HET^* and *Gli1^NULL^* vNSCs, loss of one copy of *Tgfbr2* was sufficient to increase the recruitment of *Gli1^NULL^* vNSCs, with the highest recruitment observed in mice with loss of both Gli1 and *Tgfbr2*.

To examine the fate of the recruited cells in the CC, we quantified the RFP+ cells co-expressing markers of OLs, OPCs and astrocytes. The number of OPCs generated from RFP+ cells did not alter with loss of Tgfbr2 in both Gli1^HET^ and Gli1^NULL^ vNSCs (Fig.4 F). However the proportion of OLs derived from RFP+ cells significantly increased with loss of both Gli1 and Tgfbr2 in *Gli1^NULL^;Tgfbr2^NULL^* mice, with 2.54±0.08 fold higher OLs than *Gli1^HET^;TGFβR2^HET^* mice (Fig.4G).

Taken together, these results not only indicate the synergistic effects of loss of Gli1 and TGFβ1 signaling in vNSCs, but also the role of Gpnmb in amplifying TGFβ1 signaling by inducing *Tgfbr2* leading to inhibition of recruitment and their differentiation into mature OLs.

## Discussion

Our results show that demyelination increases the levels of TGFβ1, leading to upregulation of Gpnmb expression which further activates the pathway by inducing TGFβR2 in vNSCs, ultimately blocking their differentiation into oligodendrocytes. Thus, Gpnmb is a downstream effector of TGFβ1 signaling in vNSCs and functions in a feed forward manner to sustain and amplify the pathway, resulting in inhibition of remyelination.

TGFβ1 expression is very low in the healthy brain, but is highly induced in response to injury and neurodegenerative diseases like Alzheimer’s and Parkinson’s disease, amyotrophic lateral sclerosis, stroke, traumatic brain injury and MS (Finch et al., 1993; Flanders et al., 1995; Ilzecka et al., 2002; Issazadeh et al., 1995; Krupinski et al., 1996; Lindholm et al., 1992; Mogi et al., 1995; Rimaniol et al., 1995). Likewise, our results also show that it is upregulated in the white matter CC following demyelination (Fig.3B) and induces Gpnmb in Gli1^HET^ vNSCs but not in Gli1^NULL^ vNSCs (Fig.3D). Indeed, Gpnmb expression was significantly lower in Gli1^NULL^ vNSCs compared to Gli1^HET^ vNSCs following demyelination (Fig.1A), although there was no difference in Gpnmb levels in the healthy brain (Samanta, 2021). This indicates that Gli1 is necessary for upregulation of Gpnmb following demyelination but not required for its baseline expression in healthy brains. Since TGFβ1 is known to increase Gli1 expression in various other tissues (Dennler et al., 2007), the upregulation of Gpnmb in response to demyelination is likely due to the induction of Gli1 by TGFβ1 and therefore cannot be upregulated in Gli1^NULL^ vNSCs.

Similar to TGFβ1 expression, Gpnmb is also highly expressed in many neurodegenerative diseases like amyotrophic lateral sclerosis, Alzheimer’s disease, Parkinson’s disease and MS (Hendrickx et al., 2017; Hüttenrauch et al., 2018; Moloney et al., 2018; Satoh et al., 2019; Tanaka et al., 2012). In addition to neural stem cells, Gpnmb is expressed in astrocytes, microglia, OLs and OPCs in the SVZ, CC and cortex of the healthy adult forebrain (Fig.2). However, Gpnmb is not expressed in neurons of the forebrain. In response to demyelination, the proportion of Gli1 vNSCs co-expressing Gpnmb increased significantly in the SVZ along with an increase in astrocytic expression in the CC and cortex (Fig.2), consistent with its induction by TGFβ1. Furthermore, global loss of Gpnmb significantly increased the mature OLs derived from Gli1 vNSCs without affecting the number of OPCs (Fig.1F-G), indicating its function in maturation of the oligodendrocyte lineage. Consistently, treatment of vNSCs with TGFβ1 increased the number of mature OLs, only in Gli1^NULL^ vNSCs which are unable to induce Gpnmb in response to TGFβ1 (Fig.3 E-F). This was further confirmed by directly overexpressing Gpnmb in vNSCs, which inhibited the generation of mature OLs from Gli1^NULL^ vNSCs (Fig.3 I,J).

TGFβ1 binds to the receptor TGFβR2, which is highly enriched in the neural stem cells of the adult SVZ, and inhibits their proliferation without affecting their differentiation fate into neurons, astrocytes and OPCs, but their differentiation into mature OLs had not been examined (Wachs et al., 2006). Our results show that TGFβ1→Gli1→Gpnmb signaling inhibits the differentiation into mature OLs. However, the effects of TGFβ1 are dependent on dosage and time of exposure. Indeed, we found that demyelination leads to sustained activation of TGFβ1 in vNSCs due to induction of TGFβR2 by Gpnmb, thus altering the cell fate of vNSCs (Fig.4 A,B). However, conditional ablation of TGFβR2 specifically in Gli1 vNSCs, enhanced the recruitment of both Gli1^HET^ and Gli1^NULL^ vNSC derived cells into the lesion, but increased generation of OLs only from Gli1^NULL^ vNSCs (Fig.4E-G). This points towards the presence of other Gpnmb independent mechanisms for expression of TGFβR2 in Gli1^NULL^ vNSCs, consistent with its expression in the Gpnmb^NULL^ SVZ (Fig.4B).

Taken together, these results show that enhanced TGFβ1 signaling inhibits the recruitment and differentiation of Gli1 vNSCs in demyelinated lesions in part through sustained TGFβ1 activation with a TGFβ1→Gli1→Gpnmb→ TGFβR2 feedforward loop. Identifying the cellular mediators for recruitment and differentiation in vNSCs, may ultimately provide therapeutic targets for enhancing repair and regeneration in neurodegenerative diseases.

### Limitations of study

Although we have focused on TGFβ1 because of its relevance to aging and neurodegenerative diseases, there may be other signaling pathways that regulate Gpnmb in vNSCs. We have studied the effects of global loss of Gpnmb on Gli1 vNSCs, but Gpnmb is also expressed in OPCs, astrocytes and microglia. The cell autonomous effects of Gpnmb in each cell type may be different from those observed in vNSCs. Gpnmb is a transmembrane protein that is cleaved by proteases into an intracellular domain (Gpnmb-ICD) and an ectodomain (Gpnmb-ECD). While Gpnmb-ICD can signal intrinsically by translocating to the nucleus (Utsunomiya et al., 2012), the released Gpnmb-ECD functions as an autocrine or paracrine signal by interacting with CD44 receptor in the same cell or a neighboring cell, respectively (Neal et al., 2018). In this study, we have examined the effects of complete loss of signaling via Gpnmb-ICD and Gpnmb-ECD on recruitment and differentiation of vNSCs in response to demyelination. Further studies are, therefore, needed to differentiate the cell-intrinsic signaling via Gpnmb-ICD vs. extrinsic signaling via secreted Gpnmb-ECD on remyelination.

## Acknowledgements

We thank A.R.W. and G.I. for assistance with bioinformatics analysis as well as the generation of Fig.1A and 3A. A.S.J. and M.M.T. for assistance with experiments in Fig.2, and C.A.O. with assistance with mouse colony maintenance, breeding, and cryosectioning. D.Z.R. was supported in part by a postdoctoral fellowship from the Stem Cell and Regenerative Medicine Center (SCRMC) at UW-Madison. J.S. was funded by the Marie and Jean-Pierre Boespflug Foundation for Myopathic Research and by the University of Wisconsin - Madison Office of the Vice Chancellor for Research and Graduate Education with funding from the Wisconsin Alumni Research Foundation.

## Author Contributions

Conceptualization, D.Z.R. and J.S.; Methodology, D.Z.R. and J.S.; Formal Analysis, A.W., G.I., D.Z.R. and J.S.; Investigation, D.Z.R., A.S.J. and M.M.T.; Resources, D.Z.R., C.A.O., A.W., G.I. and J.S.; Visualization, D.Z.R. and J.S.; Writing – Original Draft, D.Z.R. and J.S.; Writing – Review & Editing, D.Z.R., A.W., A.S.J., M.M.T., C.A.O, G.I. and J.S.; Funding Acquisition, J.S. and D.Z.R.; Supervision, J.S.

## MATERIALS AND METHODS

### Lead Contact

Further information and requests for resources and reagents should be directed and will be fulfilled by the lead contact Jayshree Samanta.

### Materials availability

The pLV-Gpnmb overexpression plasmid is available and will be provided by the lead contact, Jayshree Samanta upon request.

### Data and code availability

The Gli1 NSC RNAseq is available online (GSE162683), and all other GSE datasets are currently publicly available. All analysis code is available online, and no custom code was created for or from this project.

#### Fate mapping and cuprizone demyelination

All animals were used and maintained according to protocols approved by the University of Wisconsin IACUC. These mouse lines were obtained from Jackson labs or KOMP repository: Gli1^CreERT2^ (Jax# 007913), ROSA26Sor^tm9(CAG-tdTomato)Hze^ (Jax# 007909), Tgfbr2^tm1Karl^ (Jax# 012603), Gpnmb^tm1.1(KOMP)Vlcg^ (Mutant Mouse Resource & Research Center (MMRRC) #047926-UCD). The genotypes for the mouse lines are as follows: Gpnmb^HET^ = Gpnmb^tm1.1(KOMP)Vlcg/+^, Gpnmb^NULL^ = Gpnmb^tm1.1(KOMP)Vlcg/tm1.1^, Ai9 = ROSA26Sor^tm9/tm9^; Gli1^HET^= Gli1^CreERT2/+^;Ai9, Gli1^NULL^ = Gli1^CreERT2/CreERT2^;Ai9, TGFβR2^HET^ – Tgfbr2^tm1Karl/+^, TGFβR2^NULL^ – TgfbR2^tm1Karl/tm1 Karl^ . Complete genotypes and abbreviated names are summarized in Table S1. All the mice were maintained on C57Bl/6 background. 12-16 week old mice were administered 5 mg tamoxifen (Sigma) in corn oil on alternate days for a total of four intraperitoneal injections. No labelling was seen in the absence of tamoxifen administration. A week later, demyelination was induced by feeding 0.2% cuprizone in the chow for 5 weeks following which the diet was returned to normal chow for recovery from demyelination.

#### Stereotaxic Surgery

1 week prior to surgery, mice received 4 i.p. injections of 5mg tamoxifen every other day. 1% lysophosphatidylcholine (LPC) (L4129, Millipore Sigma) was dissolved in sterile saline to a final concentration of 10mg/mL. Mice were injected with 0.3mg/kg Buprenorphine 15-30min prior to induction of anesthesia, and then once every 12 hours for up to 48 hours post-surgery depending on recovery. 4-5% isoflurane with 1L/min oxygen supplementation was used to induce anesthesia until steady, rhythmic ∼60breaths/min breathing was observed, following which mice were transferred to a nose cone with a bite bar on Stoelting digital stereotaxic frame with built in heated base maintained at 38⁰C for the duration of surgery. Once immobilized in the frame, the dorsal fur of the skull was cleaned 3x with 70% ethanol, then a posterior-anterior incision was made to expose the skull. A 2mm burr hole was drilled at the following coordinates: 0.5mm AP x 1.0mm ML x -1.9mm DV (measured from the surface of the skull). A 10uL 33G Hamilton NEUROS syringe (#53497) was lowered and 1uL of 1% LPC or 1uL of sterile saline was injected over 5min. The needle was slowly retracted, the wound sealed with Vetbond (#1469SB, 3M) and mice were moved to a 37⁰C chamber to recover until they became mobile. Mice were monitored daily for recovery, and sacrificed 1 week following surgery.

#### Primary NSC culture

The brains were harvested from mice after euthanasia and placed in an acrylic mouse brain mold (BS-A-5000C, Braintree Scientific) and 1mm slices were acquired from the caudal aspect of the olfactory bulbs to the dorsal hippocampus. Brain slices were moved to dissection media composed of DMEM/F-12 (#11320032, Gibco) with 1x Antibiotic-Antimycotic (#15240112 – Gibco) and the SVZ tissue lining the ventral and lateral aspects of the lateral ventricles was microdissected from 3 consecutive slices. The dissected SVZ tissue was minced and collected by centrifugation at 400g for 5min. The tissue was then digested with 0.05% Trypsin-EDTA (#25300062, Gibco) and incubated at 37⁰C for 5min with agitation every 60-90secs to form a cell suspension. After neutralizing the Trypsin-EDTA with 1mg/mL Soybean Trypsin Inhibitor (#17075029, Gibco), the cells were needle triturated and resuspended in Mouse NSC Proliferation Media (1x Anti-Anti, 10ng/mL bFGF (#78003, Stem Cell Tech, Vancouver), 20ng/mL mEGF (#78016, Stem Cell Tech), 0.0002% Heparin Sulfate (#H7640, Sigma), 1x Mouse Proliferation Supplement (#5701) in NeuroCult Media (#5702 Stem Cell Tech). The dissociated cells were then grown as floating neurospheres in proliferation media at 37⁰C and 5% CO_2_. Primary NSCs were supplemented with fresh bFGF and mEGF at day 3 and day 5 after plating. Neurospheres were passaged at day 7 to generate secondary neurospheres. Secondary neurospheres were dissociated into single cells and plated on Matrigel (#356234 Corning, concentration tested by WiCell) coated coverslips in differentiation media (1x Anti-Anti, 1ng/mL bFGF, 2ng/mL mEGF, 0.0002% Heparin, 1x Differentiation Supplement in Neurocult Media (#5704 Stem Cell Tech Kit). Cells were maintained in Differentiation Media for 7 days with half media changes every other day, then harvested for RNA or fixed in ice-cold methanol for 15min for immunolabelling.

#### Lentivirus generation

HEK293T cells were cultured to 40% confluency in T-175 flasks in RPMI-1640 (R8758, Corning) with 10% fetal bovine serum (SH30068.03, Hyclone) and 1x Penicillin/Streptomycin (15-140-122, Gibco). Cells were transfected with either pLV-eGFP plasmid (#36083, Addgene) or pLV-Gpnmb-eGFP (custom synthesis from Synbiotech) in combination with pMD2.G (#12259, Addgene) and psPAX2 (#12260, Addgene) in equivalent molar ratios. Viafect reagent (E4981, Promega) was utilized at a 6:1 ratio for transfection as per the manufacturer’s instructions. Briefly, Viafect was added to pre-warmed serum-free media, followed by addition of all plasmids with gentle mixing and incubation at room temperature for 20min. The plasmid/Viafect mixture was added to cells and allowed to incubate for 12hours, followed by full media change. The supernatant was collected at 24 and 48 hours following the media change and pooled together before filtering through a pre-cooled 0.22□m cell filter, then pelleting via ultracentrifugation for 3hours at 75,000xg in a swinging bucket rotor. The supernatant was then aspirated, leaving a residual 50-100□l which was removed by inverting tubes onto absorbent cloth. Lentivirus pellet was then resuspended in 100□l LentiGuard reagent (Cellomics) and stored at -80⁰C until use. The average titer for all lentiviruses was ∼2×10^6^ TU/mL and was used at an MOI=0.13.

#### *In vivo* immunostaining

Mice were perfused transcardially with 4% PFA, the brains were harvested and then allowed to equilibrate for 12 hours in 30% sucrose. Tissue was then embedded and frozen in OCT media, and 20 µm coronal cryosections were obtained and processed for immunofluorescence. Slides were thawed and washed 3 times in 1xPBS for 5 min each, followed by blocking in a solution of 1xPBS/0.1%BSA/0.3% TritonX-100 with 10% normalized goat serum. Then combinations of the following antibodies: rat anti-PDGFRα (1:200, BD Biosciences), rat anti-RFP (1:1000, Chroma Tek); chicken anti-GFP (1:100, Invitrogen), mouse anti-CC1 (1:400, MilliporeSigma), anti-GFAP (1:400, Sigma), anti-MBP (1:500, Millipore) and anti-LacZ (1:2,000, Sigma) and rabbit anti-NG2 (1:200, MilliporeSigma), anti-pSmad3 (1:100 Abcam) were applied by diluting antibodies in the 1xPBS/0.1%BSA/0.3% TritonX-100 solution and incubating in a humid chamber overnight at 4⁰C. Slides were then washed in 1xPBS 3 times for 5min each, and goat anti-species secondary antibodies conjugated with Alexafluor 488, 568, or 647 (1:1,000, Invitrogen) were added to the 1xPBS/0.1%BSA/0.3%TritonX-100 solution and nuclei were simultaneously counterstained with Hoechst 33342 (1:5,000, Invitrogen). Secondary antibodies were added to sections for 1 hour, followed by 3 washes with 1xPBS for 5min each, and 1 wash with filtered water for 5min. All liquid was then removed, without allowing the slides to dry followed by coverslipping the sections with Fluoromount-G before sealing them.

#### Immunofluorescence image acquisition and analysis

Total RFP+ cells were counted using a Nikon E600 Eclipse Epifluorescent microscope with wavelength specific bandpass cubes at 20x magnification. Counting was performed by examiners blinded to the genotype and experimental condition. Epifluorescent images were obtained as Z-stacks of 5µm optical sections using a Keyence BZ-X700 microscope at 20x magnification. These images were used for single channel of total labelled cell populations in the CC, SVZ (defined as a <20□m region around the lateral ventricles), and cortex (sensorimotor cortex directly dorsal to the CC region from cingulum to cingulum). For total CC area, 4x images were obtain on Keyence for all brain slices analyzed (8-9 slices/mouse) and manually segmented for total area. For imaging of co-localized markers, 1□m optical sections were obtained on a laser scanning Leica TCS SP8 confocal microscope with LasX software. Regions of interest were stitched, merged and flattened for co-localization, and Z-stack projections were used to confirm positive cells. Results were collated in Microsoft Excel before analysis in GraphPad Prism. The investigators were blinded to allocation during image analysis and outcome assessment.

#### *In vitro* immunostaining

After 14 days of differentiation, NSC cultures were fixed by washing cells once with HBSS, then fixed with 500uL of cold methanol for 15min. Cells were grown on coverslips, which were removed from wells using fine tweezers and placed on Parafilm. This allows for reduced liquid usage as the hydrophobic Parafilm prevents liquid from leaving the coverslips. Cells were then washed 3 times with 1xPBS for 5min each, then blocked in a solution of 1xPBS with 0.1% bovine serum albumin (BSA) with 0.3% Trinton X-100 and 10% normalized goat serum for 1hour at room temperature. Following blocking, primary antibodies, chicken anti-GFP (1:100), rat anti-RFP (1:2000), rabbit anti-RFP (1:1000), rabbit anti-NG2 (1:100), rat anti-PDGFRα (1:100), mouse anti-MBP (1:100) and mouse anti-GFAP (1:1000), were added to the 1xPBS/0.1%BSA/0.3%TritonX-100 solution with combinations of primary antibodies depending on isotype and presence of GFP or RFP in cells. NSCs were incubated overnight at 4⁰C in a dark humid chamber. Coverslips were then washed 3 times with 1xPBS for 5min each, then secondary antibodies (1:1000) were added with Hoechst reagent (1:5000) for 1 hour at room temperature. Coverslips were then washed 3 times in 1xPBS followed by 1 wash with distilled water, then inverted onto a drop of Fluoromount-G on clean slides and allowed to settle overnight. Images from *in vitro* studies were acquired using Keyence BZ-X700 epifluorescent microscope from 10 fields per coverslip, across a total of 3 coverslips per marker per experimental group. Each experiment was repeated 3 times and the cell counts were averaged together. Total RFP+ (fate-mapped) cells were counted using Keyence software batch analysis counting by first counting all nuclei in the image via a threshold analysis, then confirming which nuclei were overlaid with RFP for the fate-mapped cells. This analysis was confirmed to be in >95% concordance with manual counts. Differentiated NG2+ (OPCs), MBP+ (OLs), and GFAP+ (Astrocytes) cells were counted manually using ImageJ.

#### *In vivo* qRT-PCR

For *in vivo* qRT-PCR, adult (90-120 day old) mice were sacrificed by cervical dislocation and the brain was removed and placed into the coronal 1mm mouse brain mold. 1mm sections were obtained beginning caudal to the olfactory bulbs through the dorsal hippocampus. Brain slices from 0.5mm-1.0mm relative to Bregma were identified and placed into dissection media (ice cold DMEM/F-12 with 1x Anti-Anti (#15240-062, Gibco) under a dissecting microscope. The cortex spanning -3.0mm to 3.0mm relative to midline was removed by making vertical cuts to the corpus callosum (CC), then dissecting along the dorsal aspect of the CC. CC was then removed by inserting scissors into the dorsal aspects of the lateral ventricles and cutting along the dorsal border of the lateral ventricles. Finally, SVZ was removed by dissecting around the lateral ventricles, using fiber bundles in the striatum as a guide for the limit of the SVZ tissue. Immediately following dissection, tissue was frozen in pre-cold tubes on dry ice. Total mRNA was extracted using Trizol-chloroform following the manufacturer’s instructions, and concentration and purity was determined using a NanoDrop Lite (ND-LITE-PR, Thermo Fisher). mRNA was reverse-transcribed to complementary DNA using iScript cDNA Synthesis Kit (BioRad). ITaq Universal SYBR Green Supermix (BioRad) was used to perform qPCRs in a Biorad CFX Connect thermal cycler.

#### *In vitro* qRT-PCR

Differentiated NSCs plated in 24-well plates were washed with 1x PBS, then lifted using 0.05% Trypsin-EDTA for 5min at 37⁰C. An equal volume of 1mg/mL Trypsin Inhibitor in DMEM/F-12 was added to neutralize Trypsin-EDTA, and cells were physically lifted using a cell scraper; 3 wells were pooled to generate a single technical replicate. Total mRNA was isolated using Monarch Total RNA Miniprep Kit and concentration and quality was confirmed using NanoDrop Lite. cDNA was synthesized and qPCR was run as described above.

#### Bioinformatics Analysis

For the overview of *Gpnmb* expression across samples, NCBI Gene Expression Omnibus was manually searched for Affymetrix array, bulk RNAseq or scRNAseq data sets related to mouse, rat, or human oligodendroglia, all glia, or total brain cell types. Transformed data sets were queried for reporting on *Gpnmb* expression, and if present, the rank expression value within each experimental replicate was recorded and averaged for reporting.

For bulk RNAseq analysis of *Gli1^HET^* and *Gli1^NULL^* SVZ, sample preparation, RNA collection and sequencing were performed as described in (Samanta, 2021) and the data were deposited to NCBI-GEO (GSE162683). Volcano plots were generated using the package EnhancedVolcano v1.8.0 (Blighe et al., 2020) under R v4.0.3 (R Core Team, 2020). Genes with adjusted p-values less than or equal to 0.05 and absolute values of log2 fold change greater or equal to 0.6 were considered as significant differentially expressed genes (DEGs).

The KEGG pathway analysis was done by using Advaita Bio’s iPathwayGuide (version 1902) (Ahsan and Drăghici, 2017; Donato et al., 2013; Draghici et al., 2007; Tarca et al., 2009). All measured gene expressions from the RNA sequencing data were imported into the iPathwayGuide platform, and the same thresholds of DEGs as the volcano plots were used. The iPathwayGuide utilizes the Impact Analysis method, which considers the over-representation of DEGs in the pathway and the perturbation of pathway based on the pathway topology from the KEGG database, to determine the significance of each pathway (Draghici et al., 2007; Kanehisa, 2019; Kanehisa et al., 2021; Kanehisa and Goto, 2000). The calculated p-values were then corrected for multiple tests via the Benjamini-Hochberg method (Benjamini and Hochberg, 1995) to control the false discovery rate (FDR).

#### Statistical Analysis

All results were validated by at least three independent technical and biological experiments. At least 5 sections per mouse were analyzed and data from 3–5 mice, with a mix of male and female sexes, were combined to determine the average, standard deviation, and SEM. The quantitative data were expressed as mean±SEM. Statistical analysis was performed using Student’s t test, 1-way ANOVA, or 2-way ANOVA with Tukey’s post hoc t-test. Differences were considered statistically significant at *p* < 0.05.

---

**Extended Figure 1.**
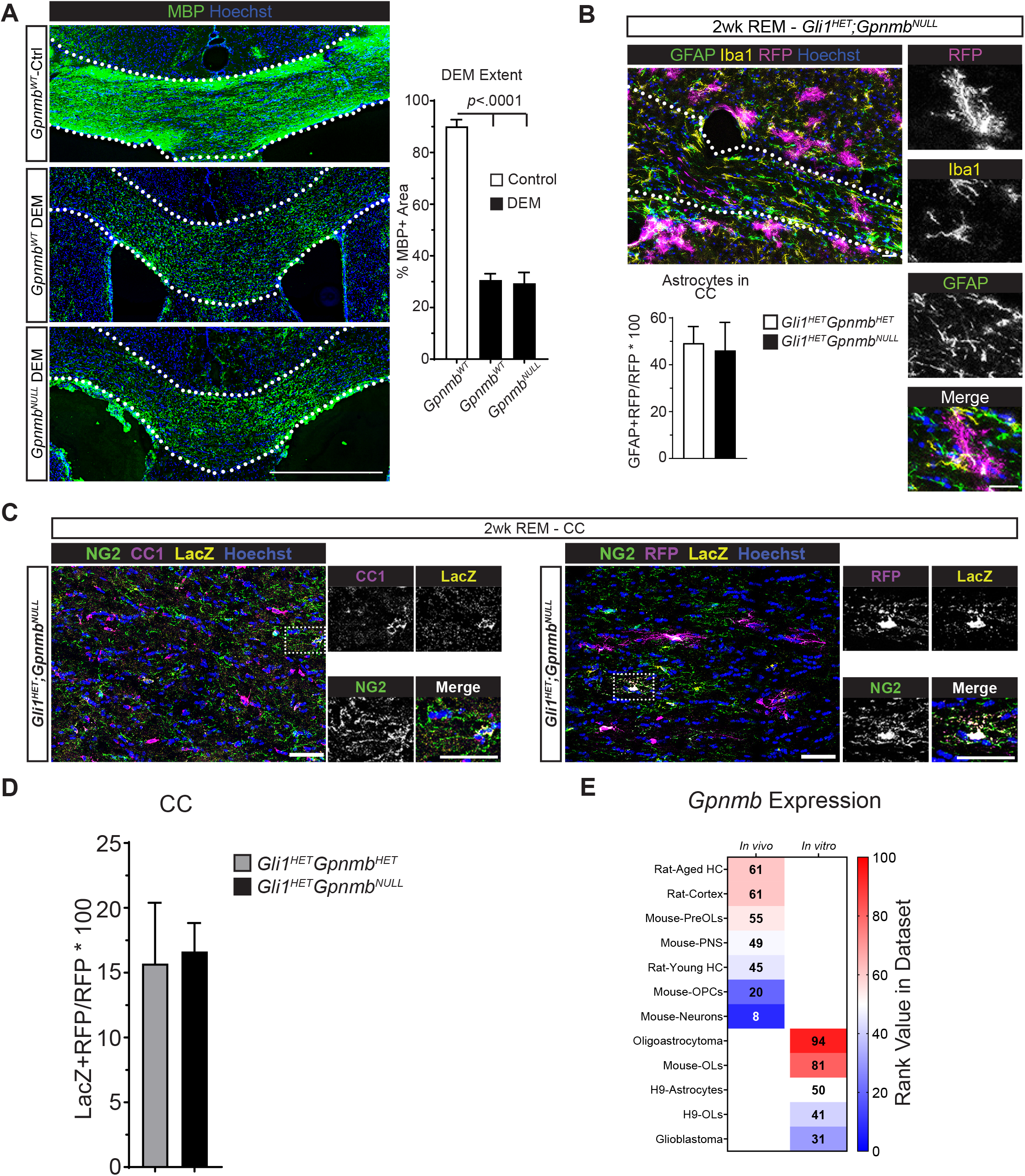
Gpnmb is expressed in multiple cell types and loss of Gpnmb does not alter extent of demyelination. (A) Immunofluorescence for MBP expression in the corpus callosum (CC) after 5 weeks of cuprizone diet (DEM) showed no difference in the MBP+ area in *Gpnmb^WT^* and *Gpnmb^NULL^* mice. n=3, scale=250µm, 1-way ANOVA with post-hoc t-test between groups. (B) Immunofluorescence of *Gli11^HET^;Gpnmb^NULL^* CC at 2 weeks of remyelination showing expression of Iba1 (yellow) and GFAP (green) along with fate-mapped Gli1 cells (RFP+). Quantification shows no change in the differentiation of RFP+ cells into astrocytes. Scale =25µm, n=3, unpaired t-test. (C) Immunofluorescence of the *Gli1^HET^;Gpnmb^NULL^* CC at 2 weeks of remyelination showing expression of LacZ in NG2+CC1 double positive cells indicating expression in pre-myelinating oligodendrocytes (left). Fate-mapped RFP+ cells in the CC co-expressing NG2, CC1, LacZ (inset). Scale = 30µm. (D) Quantification of proportion of RFP+ Gli1 fate-mapped cells co-expressing LacZ in the CC at 2 weeks of remyelination in *Gli1^HET^;Gpnmb^HET^* and *Gli1^HET^;Gpnmb^NULL^* mice. Unpaired t-tests between groups. (E) Analysis of published RNAseq and scRNAseq datasets identified *Gpnmb* expression in a variety of cells and tissues from mouse, human and rat, with lowest expression in mouse neurons.

**Extended Figure 2.**
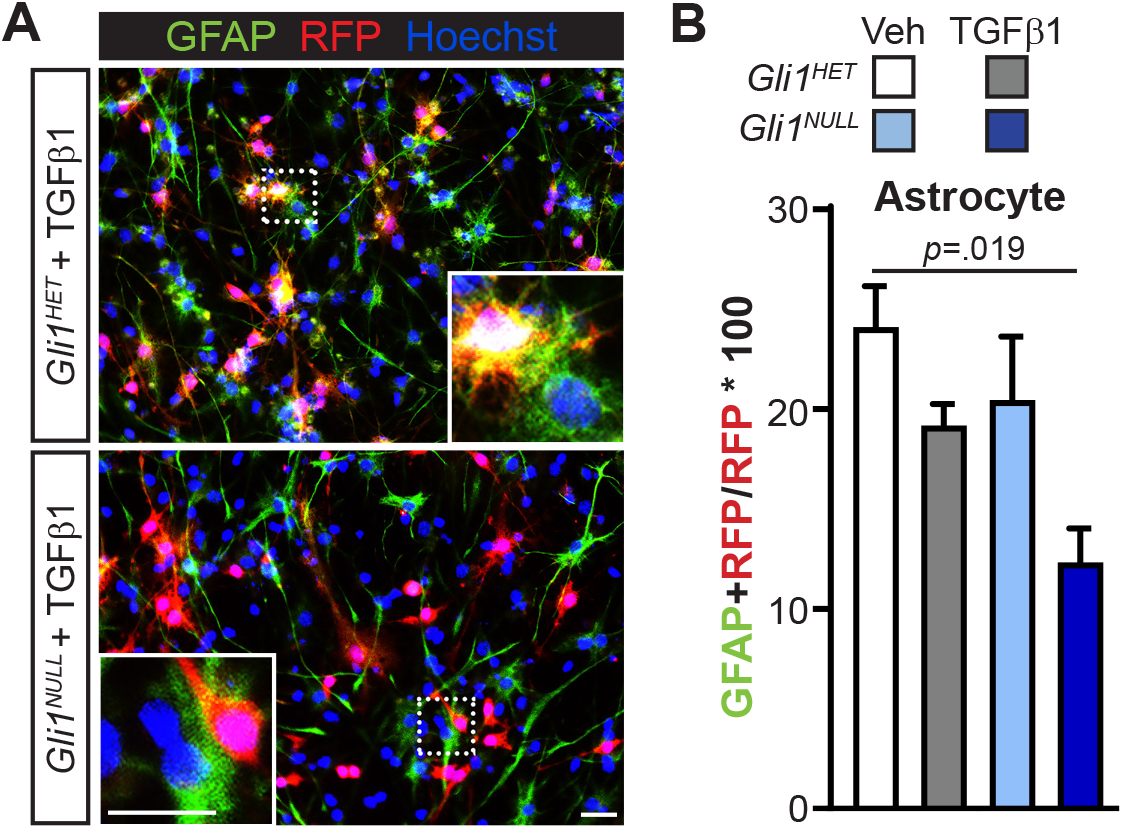
TGFβ1 does not alter differentiation of vNSC into astrocytes. (A) Immunofluorescenct images showing RFP+ vNSCs and GFAP+ astrocytes following treatment with vehicle or TGFβ1. Scale=25µm. (B) Quantification of (A) showing a significant decrease in proportion of RFP+ cells co-expressing GFAP in *Gli1^NULL^* vNSCs treated with TGFβ1 compared to *Gli1^HET^* vehicle controls. n=3, data=mean±SEM, 1-way ANOVA with post-hoc t-tests between groups.

**Extended Data 3.**
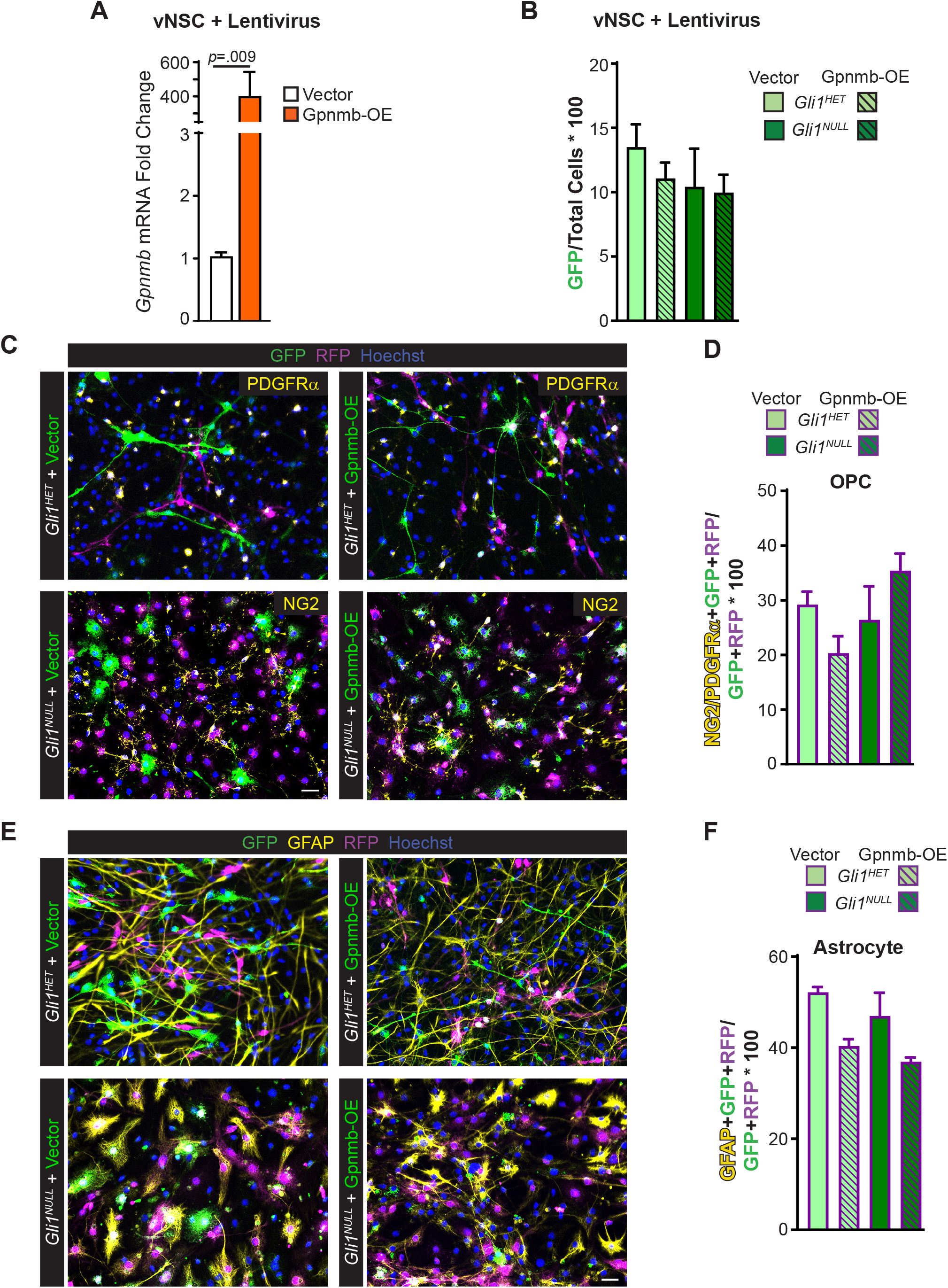
*Gpnmb* overexpression does not alter differentiation of vNSC into OPCs or astrocytes. (A) QRT-PCR for *Gpnmb* mRNA expression shows significant increase in vNSCs overexpressing Gpnmb (Gpnmb-OE) compared to empty vector. n=4, data=mean±SEM, unpaired t-test. (B) Quantification of the percentage of GFP+ lentiviral infected vNSCs shows similar rates of infection in *Gli1^HET^* and *Gli1^NULL^* vNSCs. n=3, data=mean±SEM, 1-way ANOVA with post-hoc t-tests. (C) Immunofluorescent images showing RFP+ *Gli1^HET^* (top) and *Gli1^NULL^* (bottom) vNSCs (magenta), infected with the GFP or Gpnmb-overexpression (Gpnmb-OE) lentivirus (green), co-expressing OPC markers, NG2 or PDGFRα (yellow). Scale = 25µm. (D) Quantification of (C) showing no difference in percentage of OPCs generated from the RFP+ *Gli^1HET^* and *Gli^1NULL^* vNSCs, infected with the GFP or Gpnmb-overexpression (Gpnmb-OE) lentivirus. n=3, 1-way ANOVAs within groups, followed by post-hoc t-tests for comparisons within the lentiviral treatment conditions and genotypes. (E) Immunofluorescent images showing RFP+ *Gli1^HET^* (top) and *Gli1^NULL^* (bottom) vNSCs (magenta), infected with the GFP or Gpnmb-overexpression (Gpnmb-OE) lentivirus (green), co-expressing the astrocyte marker, GFAP (yellow). Scale = 25µm. (F) Quantification of (E) showing no difference in percentage of astrocytes generated from the RFP+ *Gli1^HET^* and *Gli1^NULL^* vNSCs, infected with the GFP or Gpnmb-overexpression (Gpnmb-OE). n=3, 1-way ANOVAs within cell groups, followed by post-hoc t-tests for comparisons within the lentiviral treatment conditions and genotypes.

**Extended Data 4.**
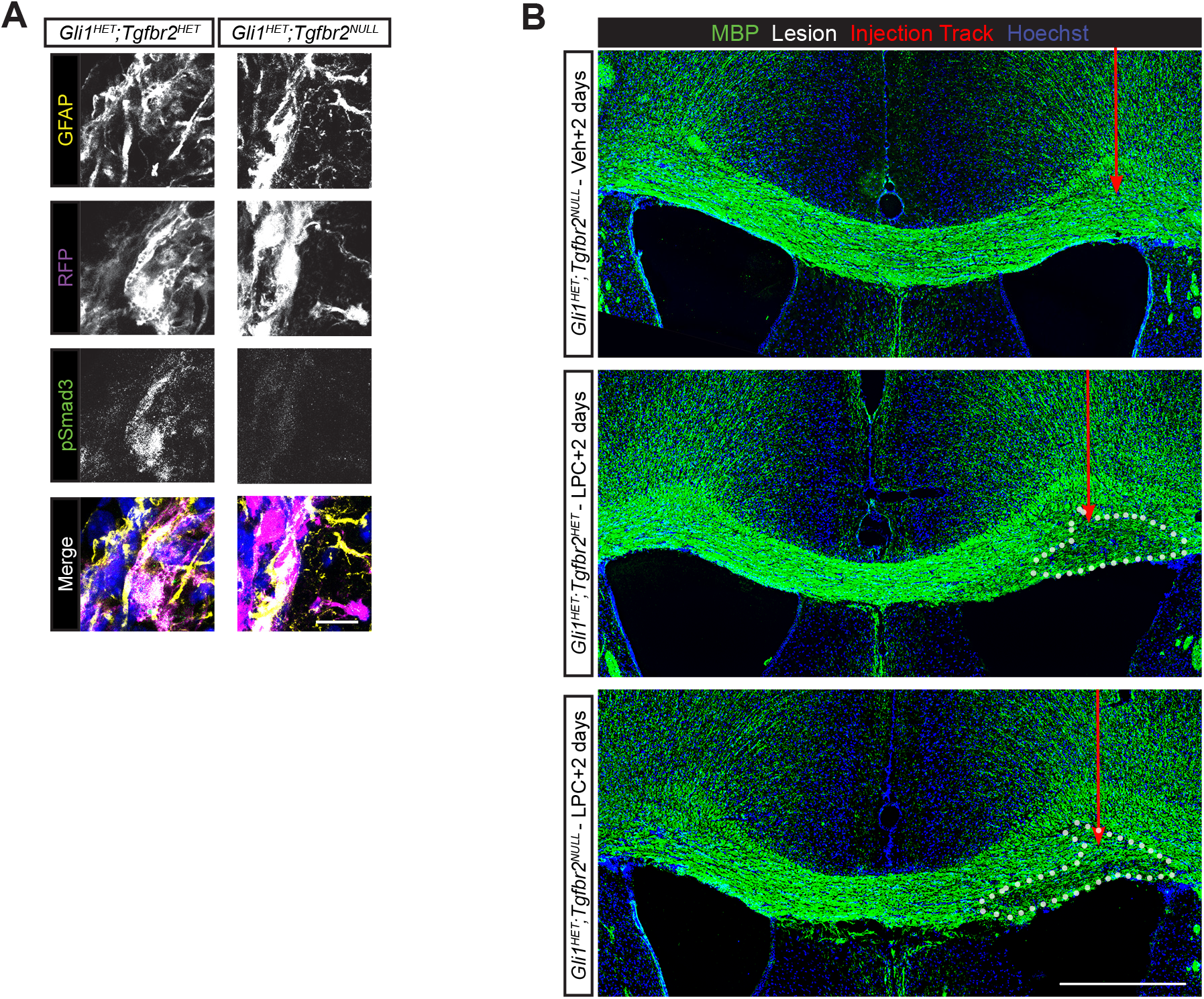
Loss of *TGFβR2* reduces pSmad3 signaling in Gli1 vNSCs and does not alter the extent of demyelination. (A) Immunofluorescent images showing GFAP+ quiescent neural stem cells (yellow), RFP+ vNSCs (magenta) and pSmad3 (green). Scale = 25µm. (B) Immunofluorescent images showing MBP+ area, 2 days after stereotaxic injection of saline or lysophosphatidyl choline (LPC) into the CC of *Gli1^HET^TGFβR2^NULL^* mice (top), *Gli1^HET^TGFβR2^HET^* (middle) and *Gli1^HET^TGFβR2^NULL^* (bottom) mice. Scale=250µm

**Supplemental Table 1.**
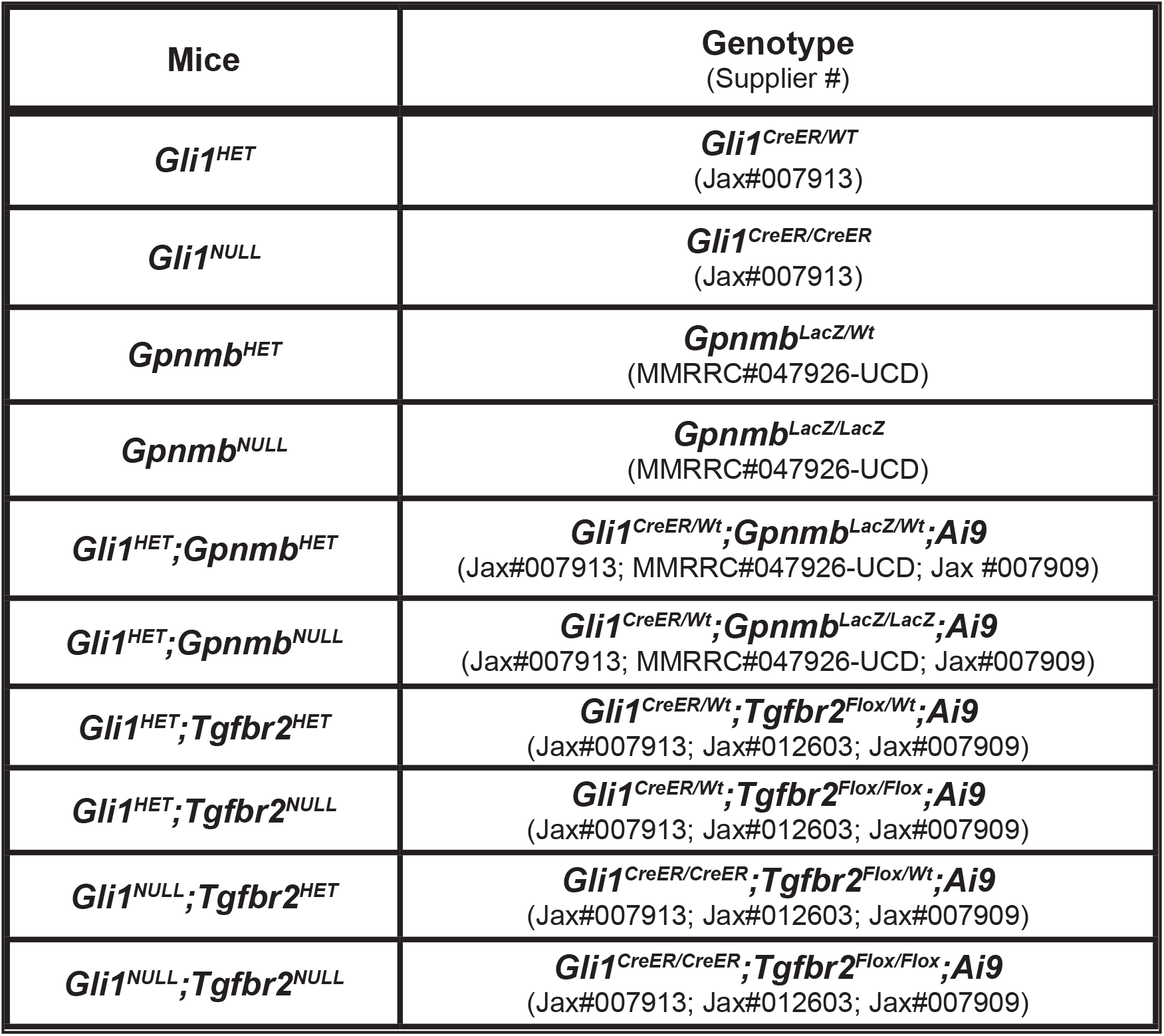
Genotypes of mice. The table lists the genotypes of all the mice, their stock numbers, and their abbreviated names.

